# Arm swapping autograft shows functional equivalency of five arms in sea stars

**DOI:** 10.1101/676254

**Authors:** Daiki Wakita, Hitoshi Aonuma, Shin Tochinai

## Abstract

Extant echinoderms show five-part radial symmetry in typical shape. However, we can find some asymmetry in their details, represented by the madreporite position not at the center, different skeletal arrangement in two of the five rays of sea urchins, and a circular cavity formed by two-end closure. We suspect the existence of any difference in hidden information between the five. In our hypothesis, deep equivalency makes no issue in function even after exchanging the position of rays; otherwise, this autograft causes some trouble in behavior or tissue formation. For this attempt, we firstly developed a method to transplant an arm tip to the counterpart of another arm in the sea star *Patiria pectinifera*. As a result, seven arms were completely implanted—four into the original positions for a control and three into different positions—with underwater surgery where we sutured with nylon thread and physically prevented nearby tube feet extending. Based on our external and internal observation, each grafted arm (*i*) gradually recovered movement coordination with the proximal body, (*ii*) regenerated its lost half as in usual distal regeneration, and (*iii*) formed no irregular intercalation filling any positional gap at the suture, no matter whether two cut arms were swapped. We here suggest a deep symmetry among the five rays of sea stars not only in morphology but also in physiology, representing an evolutionary strategy that has given equal priority to all the radial directions. Moreover, our methodological notes for grafting a mass of body in sea stars would help echinoderm research involving positional information as well as immunology.

## 1. INTRODUCTION

Extant echinoderms typically show pentaradial symmetry in apparent morphology. We recognize a regular star shape in a sea star at first glance and we collect five pairs of gonads from a sea urchin for food. The five sectors seem to have the same structure and function. Looking into it in detail, however, we become aware of some difference between them. Madreporite is a remarkable one in common echinoderms. On the aboral side of a sea star for instance, we find the madreporite as a button-like structure at a position away from the center. This is because its body has the ring canal opening outward just at one place, which is part of the water vascular system (Nichols, 1971). Thus we can identify each ray in echinoderms based on the madreporite position. One common naming is *A* to *E* clockwise in the oral view (Carpenter, 1884), where the madreporite places at the interradius between *D* and *E* in the case of general sea stars, and actually the anus also leans a bit toward the *C* to *D* region (Hotchkiss, 1995; Moore & Fell, 1966). When limited to sea urchins and some extinct taxa, another asymmetry is found in the alternate pattern of paired plates along each ambulacrum—midline area in each of the five rays. The plate series on the anticlockwise side is in advance of those on the clockwise side in three rays, whereas the other two distant rays show the opposite arrangement (Lovén, 1874). This law has been symbolized by “*aabab*” (Hotchkiss, 1978, 1995); “*a*” represents the anticlockwise-advanced ray. In the metamorphic process, echinoderms show a drastic transformation from bilaterally symmetric larva to radially symmetric adults. Here, hydrocoel—anlage of the water vascular system—initially forms a crescent, and then closes the two ends at the interradius between *C* and *D* to complete the ring (Gemmill, 1912; Hotchkiss, 1995; Mooi & David, 2008). Based on this closure position, we can postulate the first “*a*” of the “*aabab*” pattern in sea urchins is homologous to the ray *E* in sea stars (Figure 1A) although none has recognized this asymmetrical arrangement from the skeleton of sea stars (Hotchkiss, 1978, 1995). The circumoral (clockwise-anticlockwise) axis developed with exposure to spatially different expression of *Hox* genes, according to *in situ* hybridization in metamorphosing sea urchins (Arenas-Mena, Cameron, & Davidson, 2000). In behavioral terms, some studies have revealed a slight preference at the population level in moving and turning-over directions based on repeated observations of sea stars (Ji, Wu, Zhao, Wang, & Lv, 2012; Pollis & Gonor, 1975). Such cracks hidden in pentamerism would reflect an evolutionary background of echinoderms, where their early body had shown obvious asymmetry or even bilateral symmetry (Rozhnov, 2014; Sumrall & Wray, 2007).

**FIGURE 1.**
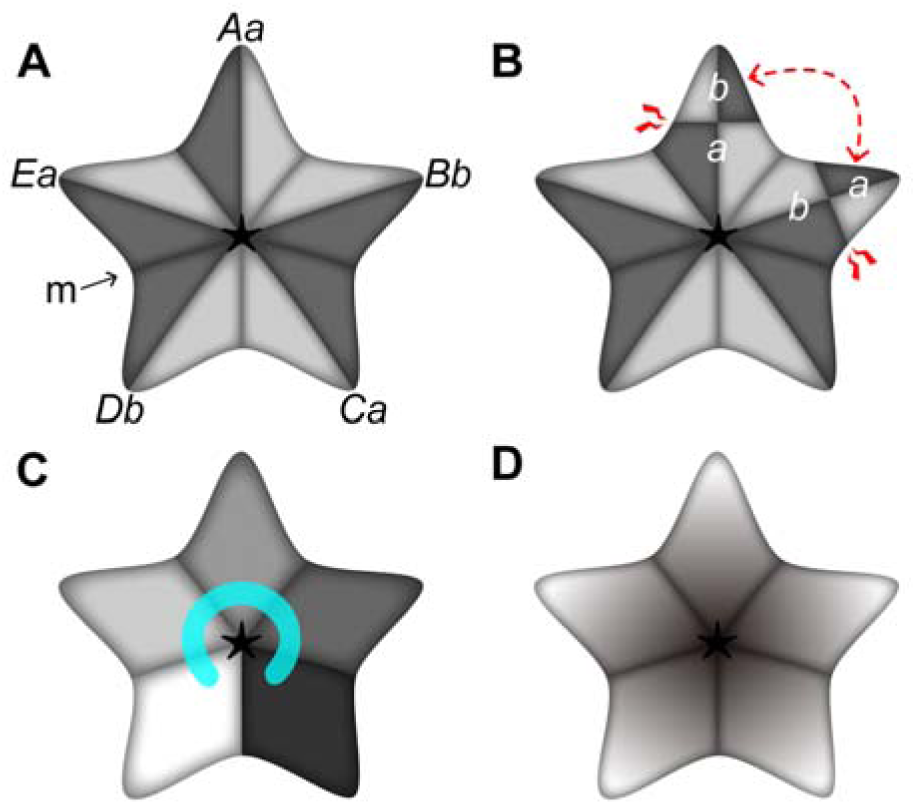
Hypotheses for positional information in the body of sea stars. **A.** Scheme of “*aabab*” pattern in the oral view. For instance, “*Aa*” denotes the ray homology “*A*” (one of Carpenter’s letters *A* to *E*) with the potential allocation “*a*” (one of the “*aabab*” pattern; the “*a*” ray shown with anticlockwise darker, the “*b*” ray shown with clockwise darker), which can be identified from the madreporite position “m.” The “*aabab*” pattern has been found in the skeleton of sea urchins but never in sea stars. **B.** Possible emergence of proximal-distal mismatches when the “*a*” and “*b*” arm tips are exchanged by autograft. **C.** Scheme where each arm has its own information. The exchange of arm tips always makes mismatches. The C shape at the center illustrates the ring canal at an early developmental stage, which makes us imagine this gradient. **D.** Scheme where all arms have the same property. No mismatch occurs through exchanging arms. Our experimental results support the last scheme

Insights in many aspects give rise to the suspicion of some potential difference among the five apparently equivalent sectors, in terms of functional expression in the adults of extant echinoderms. Our approach for this enigma is to physically exchange two sectors of the five so as to see whether some conflicts arise or not (Figure 1B), which has never been proved. We used the blue bat star *Patiria pectinifera*, in which we developed a method to transplant an arm to another arm’s position within an individual (Figure 2). In this paper, we firstly introduce autograft technique in exchanging sea stars’ arms, and then probe into any inter-arm differences based on the observation of grafts. We hypothesize the presence of asymmetry in some properties hidden in the deep structure of pentamerism. One possible gradient can be inspired by the “*aabab*” pattern (Figure 1A) in sea urchins. When the tip of a sea star’s arm potentially with the “*a*” property is grafted into the base of another arm with “*b*,” this gap would make some conflict on the suture (Figure 1B). This operation could discover the hidden existence of the “*aabab*” pattern in extant sea stars. Another simple expectation is that some property differs arm by arm, given the ontogeny where the ring canal originates from a curved linear structure (Figure 1C). In this instance, swapping autograft of two arm tips always brings proximal-distal mismatches, leading to some conflict. For a possible conflict, we suppose (*i*) impaired coordination across the suture, (*ii*) abnormal distal regeneration in the graft, or (*iii*) intercalary regeneration (Iten & Bryant, 1975) at the suture for filling a gap of positional information. The conclusion of our study is, however, contrary to the hypotheses above. We recognized no functional difference among sea stars’ five arms as schematized in Figure 1D, since arm grafts finally behaved as if they saw no exchange. Part of this work has been briefly described in conference proceedings by Wakita and Tochinai (2017), who focused on external behavior without the mention of detailed methods, failure cases, ray identification, regeneration, and internal observation; this paper details the whole.

**FIGURE 2.**
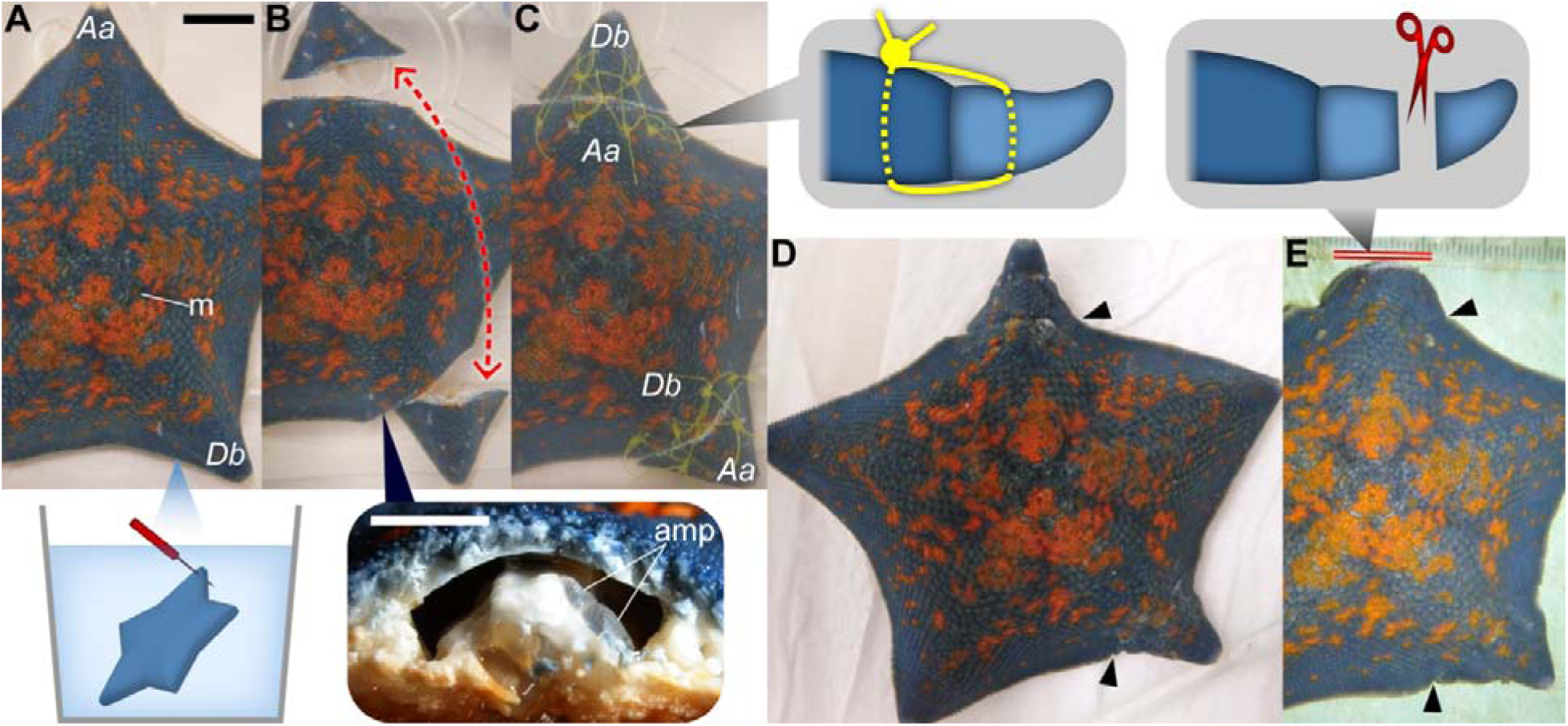
Autograft procedure where two exchanged arms were completely implanted in *Patiria pectinifera*. Photographs show the individual #27 (Table 1) in the aboral side. The grafting process A–C was performed under water throughout. **A.** The arms *Aa* and *Db* were chosen for swapping in this individual—ray identification based on the madreporite position “m”—and pierced oral-aborally at eight points for later suture (inset). **B.** The two arms were amputated at one third the radius from the tips and exchanged in position. Here, ampullae near the stump in the proximal body—“amp” in the inset—were torn to prevent their derivative tube feet actively stepping into the suture. **C.** Each exchanged arm and the proximal body were sutured with six nylon threads passing through the pre-made holes (inset). **D.** At seven days after suture, stitches were removed. Arrowheads indicate the suture position. The two whole grafts were stably implanted in this case. Later, we observed coordination and regeneration across the suture (Figures 3 and 5). **E.** At 17 days after suture, one graft was further truncated at its half (red double line and inset). The remaining base is a site for the observation of distal regeneration in the graft (Figure 4). The scale bar in A represents 1 cm, which is common in B–E while that in the B inset shows 5 mm

## 2. METHODS

### 2.1 Animals

We collected the blue bat star *Patiria pectinifera* (MÜLLER & TROSCHEL, 1842) (Asteroidea, Echinodermata; Figure 2A) at Oshoro Bay (Otaru, Hokkaido, Japan) in April to October, 2014. Sea stars were reared in laboratory aquariums in 60 × 30 × 36 cm filled with filtered aerated natural sea water at 24°C and fed with clams, shrimps, and squids once a week.

### 2.2 Autograft

For autograft experiments, we used 46 individuals which had five arms with almost equal length and had been reared in laboratory for at least a week—individuals #1–6 collected in April; #7–32 in May, and #33–46 in October; surgery conducted in April to November, 2014 (Table 1). We applied various ways to graft an arm tip into the counterpart of another arm within an individual (“Method” in Table 1). In most cases, two arms out of the five were straight amputated with scissors at one third the length of arms from the tips (Figure 2B). In several specimens, we lightly tore ampullae near the stump in the proximal body (inset in Figure 2B) with small scissors (denoted by the “torn” ampullae in Table 1). Ampulla is an accessory organ assisting the extension of its associated tube foot (Hayashi, 1935; McCurley & Kier, 1995; Nichols, 1972; Smith, 1946), so this operation was to prevent nearby tube feet interfering with later suture sites. The two distal parts were then exchanged in position without oral-aboral reversal. For a control experiment, we kept the original positions after amputation.

**TABLE 1.**
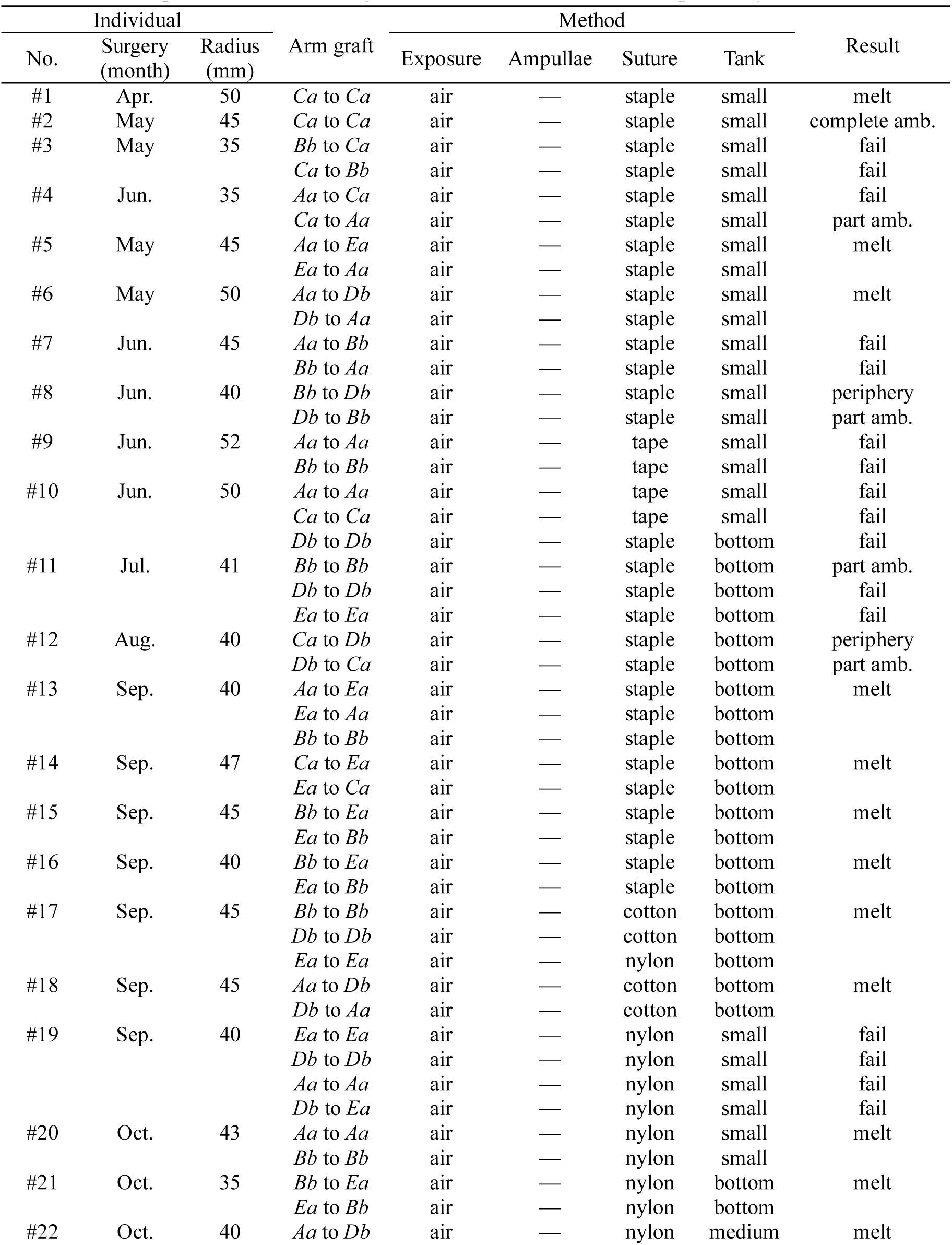
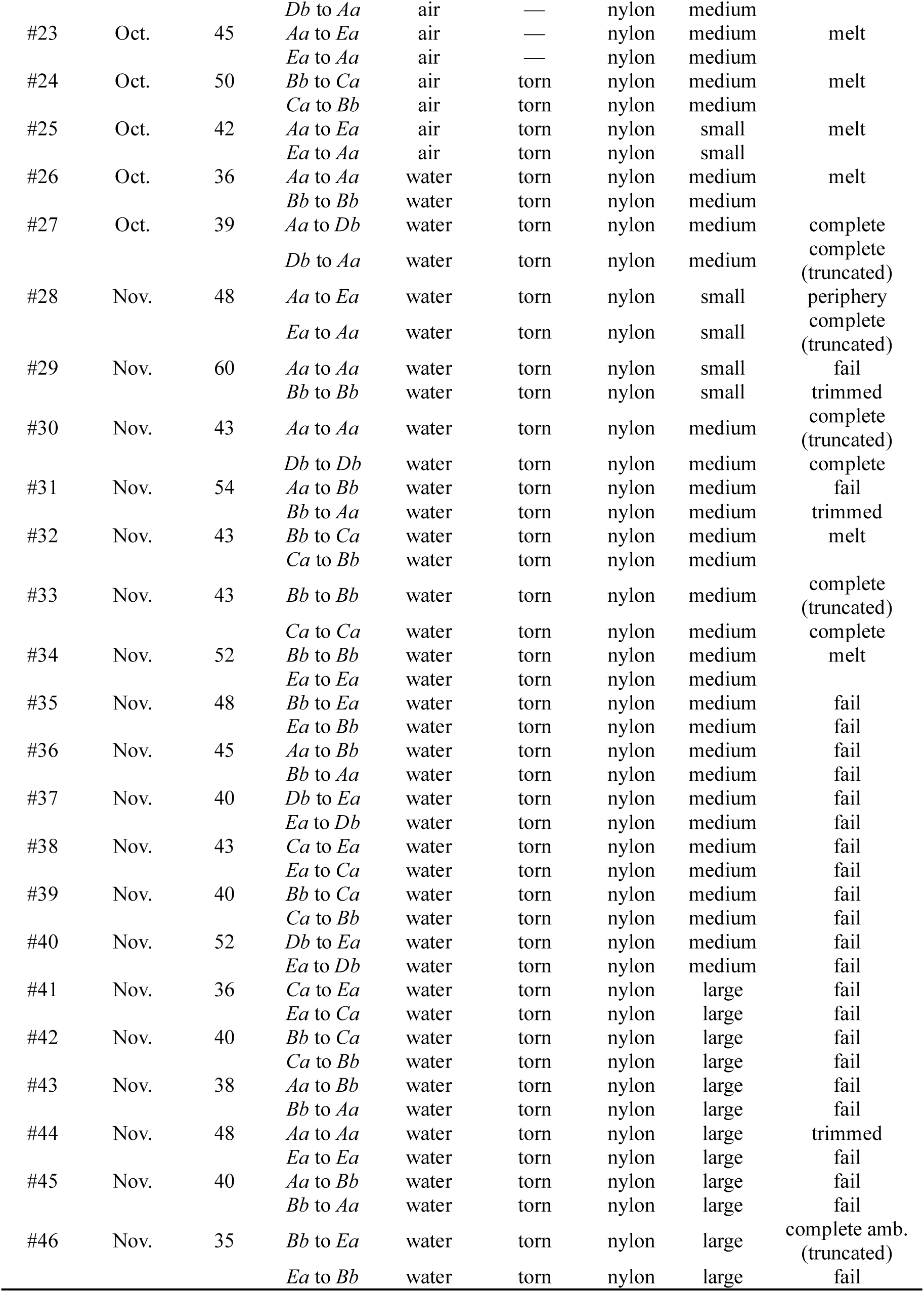
List of grafted specimens. “Arm graft” represents grafting a distal tip to a proximal base in order with lettering explained in “2.5 Terminology.” Words in “Method” and “Result” are explained in “2.2 Autograft” and “3.1 Breakdown,” respectively

Concerning how to suture the distal graft with the proximal body, we used staples (“staple” suture in Table 1; galvanized soft iron; 9.3 mm wide, 4.8 mm deep), self-fusing tape (“tape”; isobuthylene-isoprene rubber), cotton thread (“cotton”), or nylon thread (“nylon”; about 0.3 mm thick; Figure 2C). Staples were fixed from both of the oral and aboral sides, or from either side in some specimens. Self-fusing tape was tightly wrapped once about the connecting site to fit the shape of arms. When choosing cotton or nylon thread we pierced the arms at several points through the oral to aboral surfaces using a 1-mm dissecting needle, prior to amputation (inset in Figure 2A). In suture after cut, each thread was passed through the holes with a sewing needle and then both ends were knitted outside the aboral surface (inset in Figure 2C) with ligature in which we wrapped the long end around forceps three or four times and pull the short end though the coil.

Each surgery was performed with exposure to air (“air” exposure in Table 1) or strictly throughout under natural sea water without aeration (“water”; inset in Figure 2A). Post-surgery specimens were reared at 24°C in water tanks in “small” (24 × 17 × 13 cm, polycarbonate resin), “medium” (40 × 35 × 17 cm, polycarbonate resin), or “large” (60 × 30 × 36 cm, acrylic resin) size. The small and medium tanks were filled with 2.5–4.5 L of aerated natural sea water with a single individual while the large one was with more than 30 L accommodating some other individuals. For some specimens in the medium ones, we put a modified polypropylene cover of an insect cage over each (“bottom” tank in Table 1) to prevent the animal scaling the wall while burdening the suture. Sea water was changed every day for the small and medium tanks and once a month for the large one. Where the graft did not fail after one week from surgery, we removed the suture (Figure 2D) and transferred the specimen into the large tank.

### 2.3 Live observation

We observed some phenomena in living individuals where the whole or parts of grafts were stably implanted (Figure 2D). The primary focus is on movement coordination as described in the previous proceedings (Wakita & Tochinai, 2017). Coordination between grafts and proximal bodies was assessed in qualitative terms based on locomotion and food conveyance by tube feet (McCurley & Kier, 1995). Locomotion was scored in three stages when animals crept on the tank wall: almost no coordination (scarce extension and delayed detachment), poor coordination (shorter or more infrequent extension compared to the proximal one), and usual coordination. For conveyance, we gave food close to the tip of a grafted arm when the animal was stationary to observe its behavior.

Another attention was to regeneration in two aspects: whether a graft regenerates its distal end as usual; whether the suture makes any irregular intercalary regeneration. The former was testified in some grafts; after 17 days from surgery, we straight cut half the length of grafts so that the trapezoidal base of grafts remained attaching (Figure 2E; denoted by the “truncated” result in Table 1), where we observed regeneration at the stump. Hereafter, we refer to this operation as “truncation.” Indicators for the degree of regeneration were the appearance of eye spots and the extension of terminal tentacles (azygous tentacles)—measured as the ratio of the tentacle length in the grafts to the maximum tentacle length among intact arms. For comparison, we referred to normal regeneration at arm stumps after the amputation of intact arms. To address intercalation, we traced the morphology of the suture sites for a long term.

We examined coordination and regeneration per week so as to compare whether there is any difference between exchanged and non-exchanged grafts as well as matching and mismatching grafts in terms of the potential “*aabab*” pattern. Another control observation was for the behavior and regeneration of isolated arms without grafting. This supplement is to examine any difference between amputated arms with and without proximal support.

### 2.4 X-ray micro-computed tomography

We probed into how grafts make a connection with the proximal body in regard to the internal structure, using X-ray micro-computed tomography (micro-CT). Some grafted portions over the suture were trimmed with a razor and fixed in Bouin solution for about 20 months at room temperature. After fixation, samples were dehydrated with the ethanol series 70-80-90% for two days each, and stained with 1% iodine diluted in 100% ethanol for three days at 3°C to enhance X-ray reflection contrast (Metscher, 2009). They were rinsed with 100% ethanol for a day at room temperature and then moved into liquid *t*-butyl alcohol above 40°C. We put them in *t*-butyl alcohol for a day twice at 26°C, dried their surfaces on tissues for about 20 s, froze the inside *t*-butyl alcohol instantly at −20°C for 10 min to keep the original morphology, and freeze-dried them using a vacuum evaporator (PX-52, Yamato Ltd., Japan) with a cold alcohol trap (H2SO5, AS ONE, Japan). The samples were exposed to X-ray at 75 kV and 40 µA in an SMX-100CT micro-CT scanner (Shimadzu, Japan). We reconstructed and rendered the slices using VGStudio MAX ver. 2.2.6 (Volume Graphics, Germany) with the voxel size of 9–35 µm.

### 2.5 Terminology

In this paper, the letters *A* to *E* indicating ray homology are shown together with the potential “*aabab*” allocation, so to write “*Aa*,” “*Bb*,” “*Ca*,” “*Db*,” and “*Ea*” clockwise in the oral view (Figure 1A). The interradius *Ca*-*Db* accommodates the anus and the hydrocoel closure while the interradius *Db*-*Ea* contains the madreporite. In the context of autograft, the phrase “*Aa* to *Bb*” represents a site where the tip of *Aa* arm was grafted into the base of *Bb* arm, for instance.

## 3. RESULTS

### 3.1 Breakdown

All samples after autograft are summarized in Table 1. We attempted grafting 96 arms in 46 individuals in total; some individuals had more than two arms operated. Out of the whole, 18 individuals with 37 grafted arms underwent ‘melting’ of the body within two weeks after surgery, referable to as being near death. Grafts never connected in the mortal case (denoted by the “melt” result in Table 1). Among the others, where 59 arms in 28 individuals survived surgery, seven arms in four individuals were completely grafted without recognizable loss (“complete” in Table 1); three arms in three individuals were grafted with complete ambulacral regions but partial loss in periphery (“complete amb.”); four arms in four individuals were with ambulacra partly failed (“part amb.”); two arms in two individuals were with small peripheral attachments lacking ambulacra (“periphery”); three arms in three individuals were trimmed for other inquiry within a week when parts of them were more or less connected (“trimmed”); 40 arms in 21 individuals were failed in graft (“fail”).

In the exchanged case, the complete autografts were positioned at *Aa* to *Db* (distal graft to proximal body; individual #27; Figure 2), *Db* to *Aa* (#27; later truncated), and *Ea* to *Aa* (#28; later truncated). The control where we returned the amputated arms to the original places succeeded in *Aa* to *Aa* (#30; later truncated), *Bb* to *Bb* (#33; later truncated), *Ca* to *Ca* (#33), and *Db* to *Db* (#30). The partially implanted cases took place at *Aa* to *Ea* (#28), *Bb* to *Bb* (#11), *Bb* to *Db* (#8), *Bb* to *Ea* (#46; later truncated), *Ca* to *Aa* (#4), *Ca* to *Ca* (#2), *Ca* to *Db* (#12), *Db* to *Bb* (#8), and *Db* to *Ca* (#12).

### 3.2 Grafting methods

The completely grafted case in seven arms was commonly achieved by the grafting procedure shown in Figure 2. The sea star was submerged in sea water throughout surgery to prevent air passing into the body cavity (inset in Figure 2A); otherwise, we often saw post-surgery specimens swelling part of the body and eventually melting, making us suppose sealed air. Choosing two arms and imagining suture lines, we pierced each site (inset in Figure 2A) at eight points for later thread paths, so that the proximal side had four holes in a line parallel to the suture while the distal side had four holes forming a square. The two arms each were straight amputated at distally one third the radius (Figure 2B). We pricked two or three rows of ampullae near the stump in the proximal body (inset in Figure 2B); otherwise, we often observed its tube feet actively stepped into the suture or even climbed the graft to easily detach it.

We tightly sutured each site while passing six nylon threads into the holes made in advance (Figure 2C). Each thread ran from the aboral to oral sides, went across the suture, and then returned from the oral to aboral sides—one round—to make a tie with the other end (inset in Figure 2C). Each hole accommodated one or two passages of thread. In the case of suturing by staples, we never saw complete implants but seven arms were implanted with their tips or peripheries largely lost (Table 1). As to failing suture, self-fusing tape collapsed wrapped tissue or came off easily, whereas cotton thread loosened by water to miss grafts, but we employed these each only in four arm grafts.

The whole surgery using nylon thread under water took 40 to 50 min. Without exposure to air, the post-surgery individual was moved into the medium or small tank with about 3 L of sea water. The medium tank with a bottom cover and the large tank also had some partial implants (Table 1). We removed all the nylon threads after a week (Figure 2D). The implanted animals were reared in the large tank and fed as usual afterward.

### 3.3 Coordination

We observed how tube feet moved in the seven complete grafts in the individuals #27 (*Aa* to *Db*, *Db* to *Aa*), #28 (*Ea* to *Aa*), #30 (*Aa* to *Aa*, *Db* to *Db*), and #33 (*Bb* to *Bb*, *Ca* to *Ca*). In two weeks after surgery, we often saw the grafts were hung down while the proximal body normally attached to the tank wall (Figure 3A). The tube feet of the grafts poorly showed extension and adhesion to the wall (Supporting Information Video S1). When the proximal body moved in a direction, the grafts sometimes stayed in a place because some tube feet casually adhered to the substance but were difficult to detach. When a food was placed near to the graft tips, the tube feet never captured it even when the food attached. Other intact arms eventually came to catch the food to convey it proximally. These observations at early stages represent a poor ability of the grafts in coordinating locomotion and capturing food.

**FIGURE 3.**
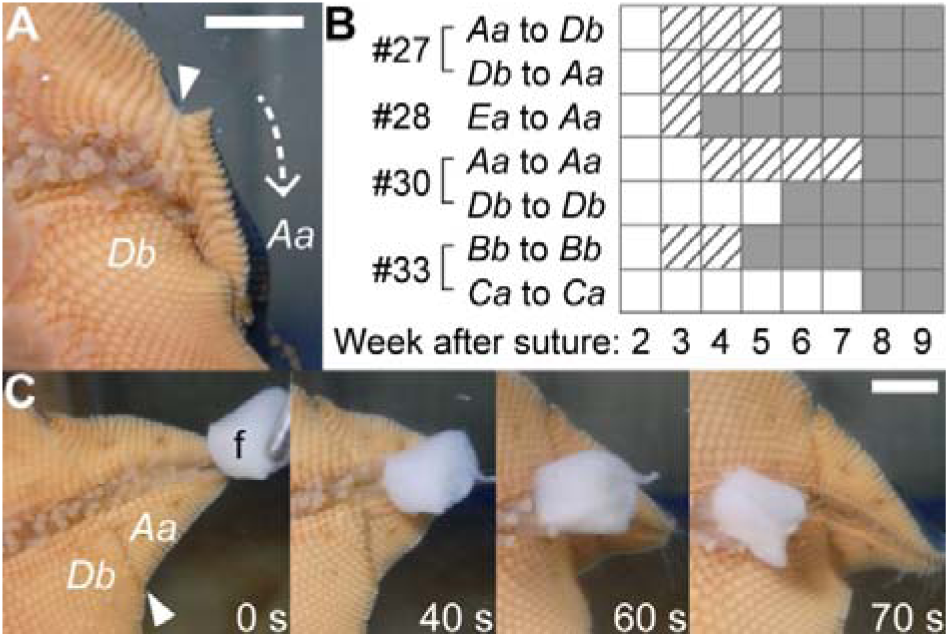
Recovery in coordination after autograft in *Patiria pectinifera*. Photographs show the *Aa* tip implanted in the *Db* base in the individual #27 (Table 1) attaching to the wall, viewed in the oral side outside the tank; arrowheads indicate the suture. **A.** Two weeks after surgery. The graft was hung down in the arrow direction and its tube feet scarcely showed active extension to the wall, contrary to the proximal ray adhering as usual. **B.** Weekly qualitative assessment of whether grafts’ tube feet show locomotion in cooperation with the proximal manner in the seven complete implants: almost no coordination shown by open squares, poor one by striped squares, and usual one by filled squares. **C.** 12 weeks after surgery. Tube feet in the *Aa* tip actively conveyed a food (“f,” squid) toward the *Db* base, as observed in normal arms. Each elapsed time is from when the tip touched the food. Scale bars represent 5 mm. Supporting Information Video S2 shows the recovery in locomotion

We found the activity of the grafts’ tube feet gradually recovered week by week (Figure 3B). By four to eight weeks after suture, we had recognized the tube feet well extended and contracted in cooperation with the proximal locomotion (Video S1). However, in feeding, other intact arms still often overrode the grafted arms at this stage. We had finally found the grafts conveying food as usual by 12 weeks after suture (Figure 3C). The timing of this recovery process more or less varied among the seven grafts, but all consistently acquired normally coordinated movement within some months. At least our results could not say the control grafts to the original positions were faster in recovery.

### 3.4 Regeneration

We observed regeneration at the tips of the five graft sites *Da* to *Aa* (individual #27), *Ea* to *Aa* (#28), *Aa* to *Aa* (#30), *Bb* to *Bb* (#33), and *Bb* to *Ea* (#46), following the truncation of each distal half at 17 days after suture (Figures 2E). These irregularly located arms, however, showed the usual process of distal regeneration (Figure 4). The truncated stumps (Figure 4A,B) closed within one day after cut. Regenerating skeletal plates became visible along the arch-shaped marginal area in two weeks (Figure 4C). Terminal tentacles were firstly recognized under water in two to four weeks (Figure 4D). In the meantime, we remarked the eye spot at the tip of each ambulacrum (Figure 4E). Later, eyespots increased the number of red ocelli while terminal tentacles increased the length of extension (Figure 4F). The common process was characterized in control stumps amputated at one third or sixth the arm length from the tips without graft, which were obtained from 14 arms in eight individuals (Figure 4G,H). In the observation timing of eye plots and terminal tentacles, the graft data lied around the average range in appearance. For the case of partial implants after exchanging arms, such as in the individuals #4 (*Ca* to *Aa*) and #8 (*Bb* to *Db*, *Db* to *Bb*), we similarly found the step-by-step regeneration of distal organs.

**FIGURE 4.**
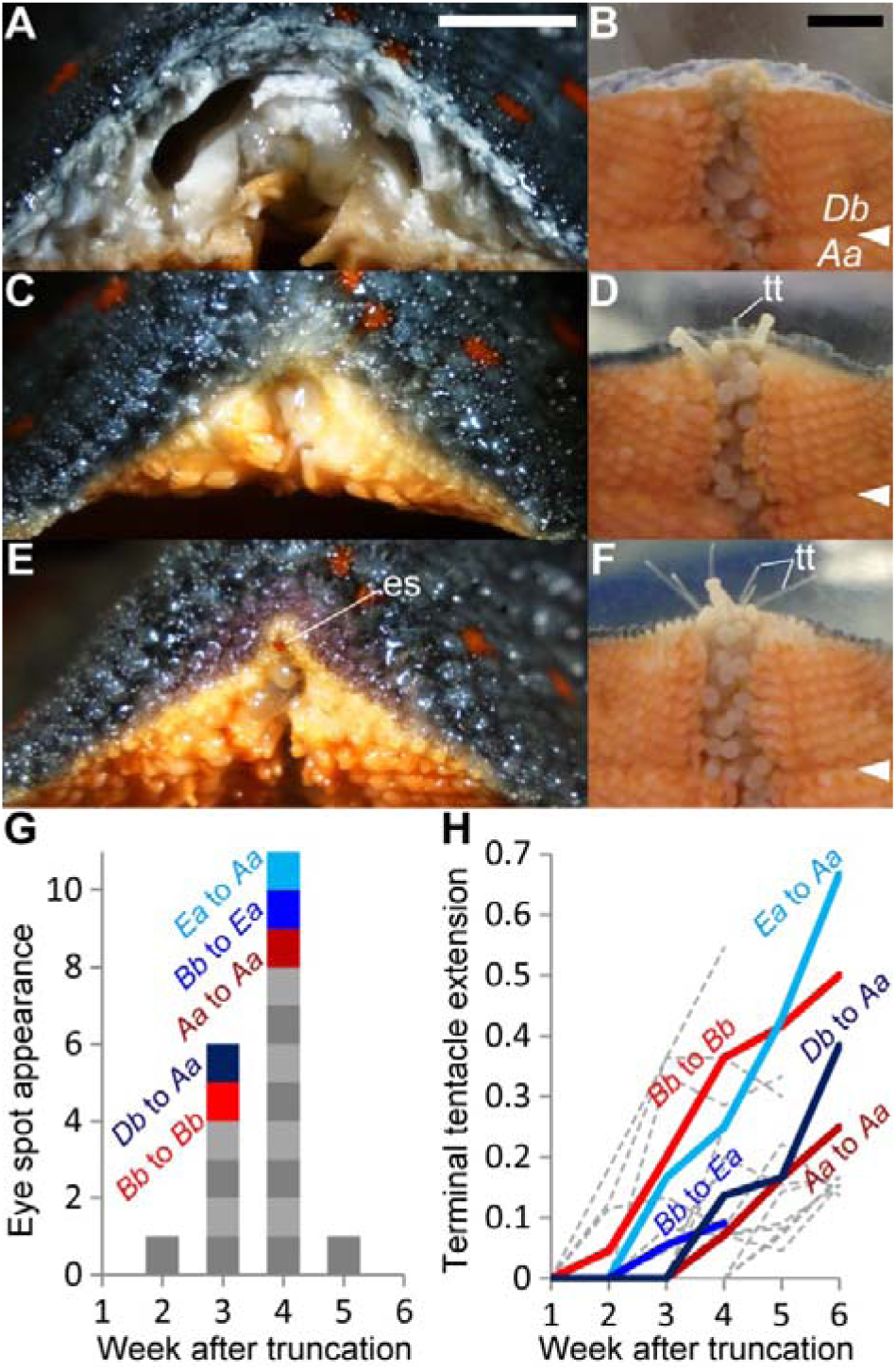
Distal regeneration in autografted arms in *Patiria pectinifera*. Photographs show the *Db* arm after truncation, with its base implanted in the *Aa* arm in the individual #27 (Table 1). **A,B.** Just after truncating the graft; 17 days after suture. **C,D.** Two weeks after truncation. **E,F.** Four weeks after truncation. **A,C,E.** Sequential images of the distal view of the stump in the air; “es” eye spot. **B,D,F.** Sequential images of the oral view under water; the sea star attaching to the wall viewed outside the tank; “tt,” terminal tentacles. Arrowheads indicate the suture. Scale bars represent 5 mm. **G.** Stacked bar chart showing when the appearance of each eye spot was found in weekly observation after truncation. **H.** Line chart showing how long terminal tentacles extended; line connecting weekly ratios—the grafts’ tentacle length to the intact arms’ maximum tentacle length. In G and H, graft sites are colored with labels (*Db* to *Aa* in the individual #27, *Ea* to *Aa* in #28, *Aa* to *Aa* in #30, *Bb* to *Bb* in #33, *Bb* to *Ea* in #46; Table 1), while gray unlabeled plots represent usual regeneration after cutting arms in non-grafted individuals as a control. The distal ends of grafts share the average regeneration process

Subsequently, the truncated grafts formed the normal distal ends, where we defined the light-colored regenerates in the aboral surface (Figure 5). At the suture lines, we never recognized irregular intercalary regeneration in all the grafted sites, while we traced them for several months or over a year after suture (e.g. #4 for 348 days; #8 for 460 days; #27 for 287 days; #28 for 109 days; #30 for 710 days; #33 for 322 days). Although each suture initially had slits at the edges and a gulf in the ambulacrum (Figure 5A), the marginal plate series and ambulacral grooves became smooth without superfluous structure over the suture (Figure 5B). The rows of oral plates were aligned one by one with the rows in the other side (shown by the dots in Figure 5A,B). In the aboral side, the very proximal verge along the suture line built some plates with light color just as in the distal regenerates, implying fresh inserts after graft (Figure 5C). Sometimes we found aboral plates lying across the line.

**FIGURE 5.**
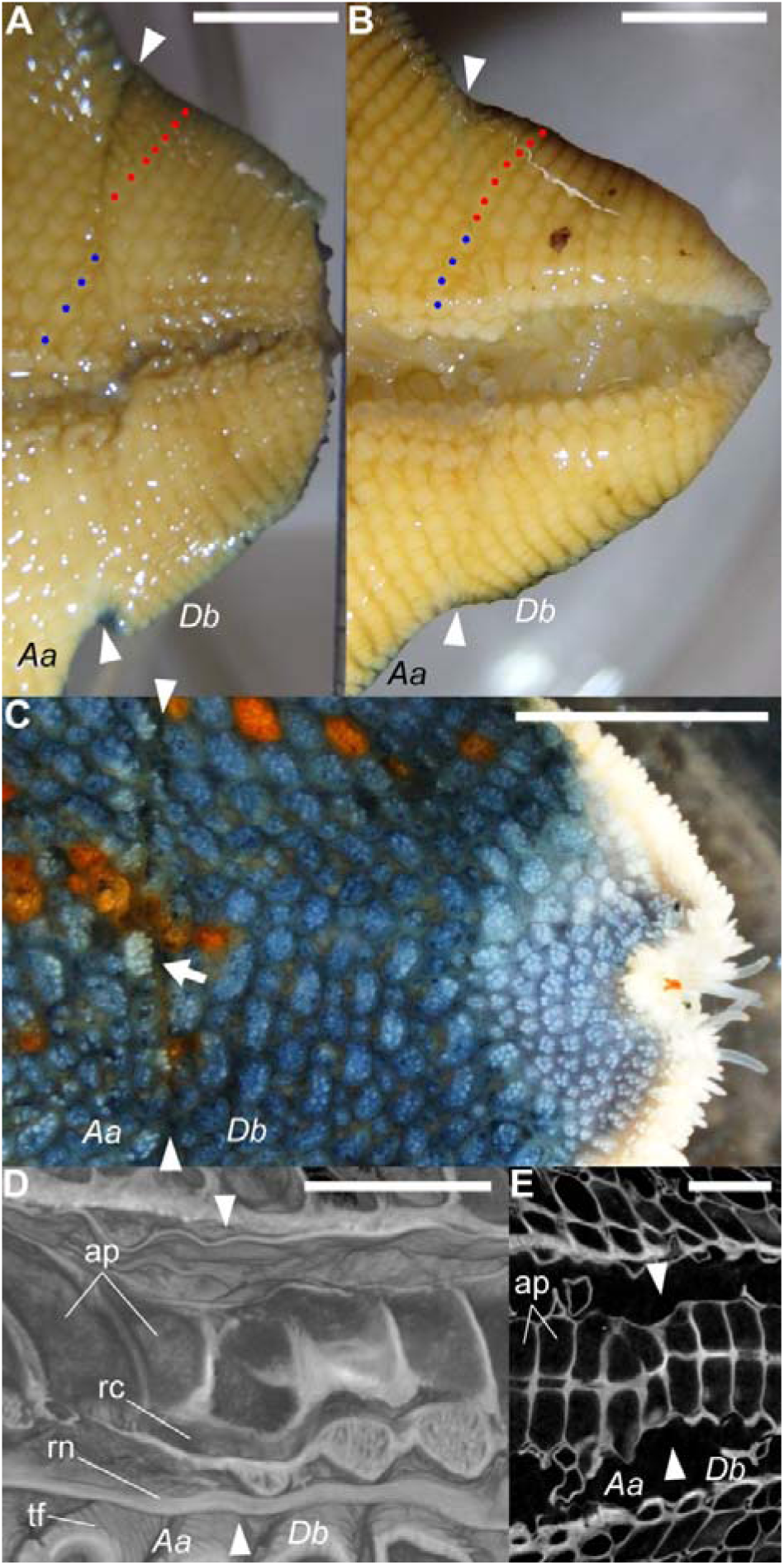
Regeneration at the suture after autograft exchanging the arms *Aa* and *Db* in *Patiria pectinifera*. Photographs show the *Db* tip implanted in the *Aa* base in the individual #27 (Table 1). Arrowheads indicate the suture and right is the distal side. **A.** 17 days after suture, just after truncating the graft; macroscopic oral view in the air. **B–E.** 41 weeks after graft; about 39 weeks after truncation. **B.** Oral view in the air. Dots represent a row of oral plates (blue in *Aa* and red in *Db*), corresponding to those shown in A. The initial gap had been reduced. **C.** Aboral view under water. The arrow at the suture indicates a light-colored plate implying fresh ones as in the regenerated tip. **D.** Three-dimensional reconstruction images of the suture obtained by X-ray micro-computed tomography (micro-CT); sliced oral-aborally along the ambulacral midline. Bottom is the oral side. Abbreviations: “ap,” ambulacral plate’s space; “rc,” radial canal, opening through the suture; “rn,” radial nerve, contained in the wall of the ambulacral groove; “tf,” tube foot. **E.** Horizontal micro-CT slice (13 µm thick) showing the arrangement of the paired series of ambulacral plates. Bottom is the clockwise direction. Scale bars represent 5 mm in A–C and 1 mm in D,E. Three-dimensional animation of these scanned images is available in Supporting Information Video S3 (for other graft sites, see Supporting Information Videos S2, S4, and S5)

Such continuity could also apply to the internal or microscopic structure imaged by micro-CT scanning (Figure 5D,E)—we scanned the graft sites of *Aa* to *Db* (individual #27; Supporting Information Video S2 for three-dimensional animation), *Db* to *Aa* (#27; Supporting Information Video S3), *Ea* to *Aa* (#28; Supporting Information Video S4), and *Bb* to *Bb* (#33; Supporting Information Video S5). In all the imaged samples, the outer lines of ambulacral grooves were smoothly connected so that it was difficult to define the suture positions here (Figure 5D). Between these surfaces and ambulacral plates inside, we recognized radial canals, which shared the opening with ampullae and tube feet, to be open continuously across the suture (easily traceable in Videos S2–S5 in the last distal-proximal slicing animations) even where it was considerably dislocated in the lateral direction (conspicuous in #28’s *Ea* to *Aa*; Video S4). The series of ambulacral plates were also connected while more or less curving at the suture’s proximal side (Figure 5E). The plate series on the clockwise and anticlockwise sides were paired throughout, except for the site *Aa* to *Db* in #27, where one ambulacral plate was azygos at the suture (Video S2).

### 3.5 Isolated arms

We observed nine arm tips amputated from five individuals at one third the length of arms. The isolated arms showed locomotion and sometimes scaled the wall. When we put a food closely to their tips, they more or less extended terminal tentacles but never attempted to creep toward or convey it even when the food attached. The isolates did not melt for at least five days; the longest recorded 11 weeks. In the experiment that we further truncated isolated arms—three obtained from three individuals—, the basal trapezoidal halves melted within a week, so we were unable to observe their distal regeneration.

## 4. DISCUSSION

### 4.1 Transplantation in sea stars

We described an autograft method in sea stars where seven arms were practically exchanged without recognizable loss (Figure 2). Here, we performed whole surgery underwater, pierced later suture sites for thread, amputated two arm tips each at one third the radius, tore ampullae nearby the proximal stumps, sutured with nylon thread, and removed the stitches after a week. Development of the grafting technique in this animal would expand not only the accessibility to potential asymmetry in the bizarre body plan but also the breadth of immunology in echinoderms. As reviewed by Smith et al. (2018), many researchers have paid attention to echinoderms’ immune system in the aspects of, for example, ecologically influential mass mortality, genomics in the purple sea urchin *Strongylocentrotus purpuratus*, and similarity to vertebrates as a deuterostome. Our study would be part of this field with respect to a methodological contribution.

As to previous grafting experiments in sea stars, King (1898) performed allograft in an *Asterias* species, where two individuals were cut across the disk and one two-armed portion was tied to the other individual’s three-armed portion. This attempt obtained the success rate of one out of 72 using course thread—the successful chimera made the suture ectoderm continuous in about two weeks and lived for three weeks. Although how he made a tie is not described, we can see a common difficulty in grafting a mass of body in sea stars. In terms of transplantation in a smaller tissue of the body wall, other studies have demonstrated allograft is rejected while autograft is accepted in a *Dermasterias* sea star (Hildemann & Dix, 1972; Karp & Hildemann, 1976). Our low success rate in such acceptable autograft would be primarily caused by that the graft site directly borders the highly mobile series of proximal tube feet. As one solution made in our study, tube feet’s invasion into the suture was prevented by pricking their ampullae in advance during surgery (Figure 2B), in consideration of ampulla’s work to support the elongation of its linking tube foot (McCurley & Kier, 1995; Nichols, 1972; Smith, 1946). Rearing at lower temperature might be another way to reduce the entire mobility, whereas we conducted it only at room temperature. Additional difficulty would come from the peculiarity of echinoderm connective tissue to drastically change its stiffness (Motokawa, 1984). As far as we observed, sea stars softened the body for daily flexible movements, which made the suture loosen to much decrease the stability of grafts. Some treat to inhibit the change in stiffness might increase the success rate; acetylcholine and some neuropeptides are known to harden the tissue (Birenheide et al., 1998; Motokawa, 1981) while we should consider their effect to implantation.

Among 16 arms which underwent complete or partial implantation, 12 arms were exchanged as pairs in six individuals (Table 1). This bias implies the success of arm implantation would be affected more by individual properties rather than arm-by-arm surgical conditions. In this context, the continuous occurrence of melting in the individuals #13 to #26 but #19 might be partly caused by intrinsic factors in animals. Possible one is a seasonal factor as these were operated in September through October, overlapping with the breeding season of *Patiria pectinifera* reported from near the collect site (Takahashi, 1979).

In the meantime, the strictly underwater surgery would increase the survival rate given only two melting individuals out of 21, contrary to 15 melting ones out of 25 which underwent exposure to air (Table 1). We suppose suture in the air easily enclosed air bubbles in the body cavity to cause harm. Rearing environment after surgery would be of less importance since each yielded some implanted arms. As to the other major attempt in suture, staples brought not complete but partial implants in several arms. One reason for the distal or peripheral loss of grafts seems to be that oral and aboral staples made different penetrations from each surface, resulting in excessive injury for the flat arms of *P*. *pectinifera*. This rapid suture might be useful in species with a thicker body.

### 4.2 Proximal-distal movement coordination

We hypothesized arm-by-arm difference in function would bring impaired coordination between the distal graft and the proximal body after swapping two arms. Although we saw such disharmony in a few weeks after surgery (Figure 3A), the grafts showed gradual recovery in external behavior (Figure 3B,C; Video S1), regardless of whether an amputated arm was transferred to another arm position or returned to the original. This answer reflects that different arms indeed have function replaceable to each other.

The tip of an arm itself has no ability to convey food proximally, according to the observations that isolated arms never showed this attempt and grafts just after surgery exhibited a similarly poor behavior. We can suppose the conveyance function in tube feet was retrieved by a certain connection to the proximal body. Crucial ones are likely to be a radial nerve cord and a radial canal running along each arm, given the nervous system and the water vascular system both drive tube feet (Cobb, 1967; Hayashi, 1935; McCurley & Kier, 1995; Smith, 1946). Radial nerves run between ambulacral epidermis and coelomic epithelium (Cobb, 1967; Smith, 1937, 1946), so neural reconnection at the suture is highly probable based on the scanned images showing a significant continuity in the lining of ambulacral grooves (Figure 5D; Videos S2–S5). Radial canals were indeed recognized as a continuous opening over the suture, evidencing water communication between the grafts and the proximal body. This explanation can also apply to the case of locomotion by tube feet where the grafts apparently recovered the proximal-distal coordination (Video S1). Our key implication here is that an arm of sea stars integrates its behavior even where a driving network is partly made by the cells of a different arm.

### 4.3 Distal regeneration and cell migration

Our results for distal regeneration after graft truncation (Figures 4 and 5A–C) were contrary to our first expectation that an arm graft sited on another arm would have a poor ability to regenerate its tissues. Molecular marking of cell proliferation by Hernroth et al. (2010) indicated that cells constituting a regenerating arm are recruited from organs distant to the regenerate in sea stars’ body. This previous work suggests the regenerating tips of grafts were derived from the proximal body over the suture. The hard survival of truncated isolates also upholds the importance of proximal support for the maintenance of grafts. As far as we observed, the truncated ends of grafts exhibited a normal regeneration which agrees with our non-graft cut experiment as well as other reports on arm cut in sea stars (Khadra et al., 2017; Mladenov, Bisgrove, Asotra, & Burke, 1989). Thus, this consequence can represent no issue in transferring cells through a pathway containing a different arm.

### 4.4 Positional information along the circumoral axis

What the suture site eventually came to a usual form as if we made no swap (Figure 5) reinforces the deep equivalency among the five arms. We initially presumed some irregular structure would take shape at the boundary where an arm base is adjacent to a different arm tip. This idea was inspired by autograft in other animals, which has generalized that a gap of positional information over the suture makes “intercalary regeneration” to smoothly compensate the gap. This phenomenon is represented by swapped fingers in newt (anterior-posterior and proximal-distal gaps; Iten & Bryant, 1975), truncated legs in cockroach (proximal-distal gap; French, 1976), and the translocated head in planaria (anterior-posterior gap; Agata, Tanaka, Kobayashi, Kato, & Saitoh, 2003). In the instance of newt, a suture site where the left forelimb is autografted to the counterpart of the right forelimb has an anterior-posterior mismatch, so two right-forelimb-like supernumeraries often develop to make anterior-posterior gradients continuous (Iten & Bryant, 1975). With this in view, the autograft made in our study is regarded as to testify some circumoral (clockwise-anticlockwise) gradient in the body or each arm of sea stars. The relationship between the “*a*” and “*b*” arms could be rather similar to that between left and right limbs in newt. If they hide the “*aabab*” pattern in a molecular level (Figure 1A), the “*a*” to “*b*” suture would intercalate supernumerary ambulacra—radial nerves, radial canals, series of tube feet, etc.—as each ambulacrum lies as a boundary structure between the potential “*a*” and “*b*” sides in the normal body. Under the expectation that some arm-by-arm property presents (Figure 1C), the “*A*” to “*D*” suture might build an extra “*E*”-being structure.

Against these hypotheses, our case study realized nine arms in five individuals which made no irregularly large intercalation in external and internal structures for several months or even a year after grafting an arm to another (Figure 5; Videos S2–S5). Here, four arms in two individuals simultaneously had a mismatch in the potential “*aabab*” pattern (c.f. Figures 1B and 2). The proximal-distal contact in the early post-graft period probably influences the form of later connection. In many cases, the suture sites had lateral dislocation to some degree. Despite this initial bother, we saw sea stars’ attempt to adjust it using existing structure as much as possible. Remarkably, each row of oral plates regularly bridged the potential “*a*” and “*b*” areas after exchanging the arms *Aa* and *Db*, as if two incomplete lines each met a suitable partner (Figure 5A,B). The pairs of ambulacral plates—major axial skeleton in sea stars as well as other echinoderm taxa (Mooi & David, 2000)—also signified such preferred approach to the other side of the gapped suture (Figure 5E). Even where the suture had a large dislocation as in #28’s *Ea* to *Aa* site (Video S4), radial canals on the two sides succeeded in reopening one tube in a snaky way (Figure 5D). These experimental results indicate that cells in each organ recognized another arms’ counterpart as something equal that can neighbor each other. We can therefore suggest sea stars are in pentaradial symmetry even in molecular properties, as schematized in Figure 1D.

### 4.5 Conclusion

To sum up our study on sea stars, swapping two arms—even striding across the potential “*aabab*” pattern—brought no recognizable conflict in movement coordination, cell migration, and positional information. What a ray works even in part of another ray makes a suggestion of functional equivalency between five rays in sea stars. This outcome gives prominence to the high degree of their pentaradial symmetry, not limiting in morphology but extending to physiology. In particular, we reject the existence of the sea star “*aabab*” pattern by a different means from the traditional one which has merely focused on stationary shapes. Given that fossil records have implied the asymmetrical origin of the current pentamerism in echinoderms (Rozhnov, 2012; Sumrall & Wray, 2007), our paper depicts an evolutionary strategy in sea stars that has extremely reduced ray-by-ray difference in ‘active’ terms so as to equalize duties in all the radial directions.

## Acknowledgments

Our work is supported by JST CREST (grant number JPMJCR14D5) partly in micro-CT scanning.

## Supporting Information

**Video S1** Recovery in coordinated locomotion at the graft site *Aa* to *Db* in the individual #27

**Video S2** Three-dimensional animation of the graft site *Aa* to *Db* at 41 weeks after suture in the individual #27 (voxel size 34 and 18 µm)

**Video S3** Three-dimensional animation of the graft site *Db* to *Aa* and its distal regenerate at 41 weeks after suture in the individual #27 (voxel size 32 and 9 µm)

**Video S4** Three-dimensional animation of the graft site *Ea* to *Aa* at 16 weeks after suture in the individual #28 (voxel size 35 and 15 µm)

**Video S5** Three-dimensional animation of the graft site *Bb* to *Bb* at 46 weeks after suture in the individual #33 (voxel size 26 and 15 µm)

